# ZFP207 *controls pluripotency by multiple post-transcriptional mechanisms*

**DOI:** 10.1101/2021.03.02.433507

**Authors:** Sandhya Malla, Devi Prasad Bhattarai, Dario Melguizo-Sanchis, Ionut Atanasoai, Paula Groza, Ángel-Carlos Román, Dandan Zhu, Dung-Fang Lee, Claudia Kutter, Francesca Aguilo

**Affiliations:** Department of Medical Biosciences, Umeå University, SE-901 85, Umeå, Sweden; Department of Molecular Biology, Umeå University, SE-901 85, Umeå, Sweden; Wallenberg Centre for Molecular Medicine, Umeå University, SE-901 85, Umeå, Sweden; Department of Microbiology, Tumor and Cell Biology, Karolinska Institute, Science for Life Laboratory, SE-171 77, Stockholm, Sweden; Department of Biochemistry, Molecular Biology and Genetics, University of Extremadura, 06071, Badajoz, Spain; Department of Integrative Biology and Pharmacology, McGovern Medical School, The University of Texas Health Science Center at Houston, Houston, TX 77030, USA; Center for Precision Health, School of Biomedical Informatics, The University of Texas Health Science Center at Houston, Houston, TX 77030, USA; The University of Texas MD Anderson Cancer Center UTHealth Graduate School of Biomedical Sciences, Houston, TX 77030, USA; Center for Stem Cell and Regenerative Medicine, The Brown Foundation Institute of Molecular Medicine for the Prevention of Human Diseases, The University of Texas Health Science Center at Houston, Houston, TX 77030, USA

## Abstract

The pluripotent state is not solely governed by the action of the core transcription factors Oct4, Sox2, and Nanog, but also by a series of co-transcriptional and post-transcriptional events, including alternative splicing (AS) and the interaction of RNA-binding proteins (RBPs) with defined subpopulations of RNAs. Zinc Finger Protein 207 (ZFP207) is an essential transcription factor for mammalian embryonic development. Here, we employ multiple functional analyses to characterize the role of ZFP207 in mouse embryonic stem cells (ESCs). We find that ZFP207 plays a pivotal role in ESC maintenance, and silencing of *Zfp207* leads to severe neuroectodermal differentiation defects. In striking contrast to human ESCs, ZFP207 does not transcriptionally regulate stem cell and neuronal-related genes but exerts its effects by control AS networks and acting as an RBP. Our study expands the role of ZFP207 to maintain ESC identity, and underscores ZFP207 functional versatility with key roles in neural fate commitment.

## Introduction

Mouse embryonic stem cells (ESCs) are derived from the inner cell mass of the pre-implantation blastocyst. These cells exhibit unlimited self-renewal capacity and, under appropriated stimulus, retain the potential to differentiate into the three germ layers (Bradley et al., 1984). Mouse ESCs are a useful model to study early mammalian development as their differentiation potential is more robust than that of ESC-like cells derived from other mammals such as humans, which exhibit primed pluripotency and represent a more advanced embryonic stage (Ginis et al., 2004; Nichols and Smith, 2009).

Zinc finger-containing proteins (ZFN or ZFP for human or mice, respectively) are among the largest family of proteins, commonly containing a minimum of one zinc-finger (ZnF) domain which recognize DNA sequences with high affinity. This family of transcription factors play important roles in a variety of cellular processes including development, cellular differentiation, metabolism and oncogenesis (Cassandri et al., 2017). Although ZNF/ZFPs were initially classified as transcription factors, several studies have highlighted additional functions of ZNFs. For instance, it has been shown that ZFP217 could recruit the methyltransferase-like 3 (METTL3) into an inactive complex and hence, restrict N6-methyladenosine (m^6^A) deposition at pluripotency transcripts (Aguilo et al., 2015; Lee et al., 2016). In addition, recent studies identified that ZnF domains can bind RNA, and many ZNF/ZFPs act as putative RNA-binding proteins (RBPs) (Brannan et al., 2016). Indeed, analysis of quantitative global mRNA-protein interaction approaches identified ZNF207 (also termed BuGZ; Bub3 interacting GLEBS and Zinc finger domain-containing protein) as an RBP, among other ZNFs (Baltz et al., 2012; Castello et al., 2012).

ZNF207 is conserved in eukaryotes. It associates with Bub3 and with spindle microtubules to regulate chromosome alignment (Jiang et al., 2014; Jiang et al., 2015; Toledo et al., 2014). Furthermore, both ZNF207 and Bub3 interact with the spliceosome and are required for interphase RNA splicing (Wan et al., 2015), yet its specific molecular role remains elusive.

In human ESCs, ZNF207 functions as a critical transcription factor by transcriptionally regulating the expression of the pluripotency factor OCT4 (Fang et al., 2018), thereby being implicated in the maintenance of self-renewal and pluripotency. Likewise, ZNF207 has been shown to enhance reprogramming efficiency towards pluripotency (Toh et al., 2016). Alternative splicing (AS), in which splice sites in primary transcripts are differentially selected to produce structurally and functionally distinct mRNAs, plays a critical role in cell fate transitions, development, and disease (Gabut et al., 2011). ZNF207 undergoes AS during somatic cell reprogramming and differentiation of human ESCs, an isoform switch that seems to be required for the generation of induced pluripotent stem cells (iPSC), and it might also be necessary to maintain ESC self-renewal and to induce proper differentiation programs (Fang et al., 2018; Toh et al., 2016).

Here we show that ZFP207 plays an important role in the control of mouse ESC identity by a distinct mechanism, which differs from the one observed in human ESCs. Specifically, in mouse ESCs, ZFP207 does not regulate *Oct4* transcription but increases OCT4 protein stability by disrupting ubiquitin-dependent proteasomal degradation. In addition, depletion of *Zfp207* results in pluripotency defects and blocks neuroectodermal specification without significant changes in the transcriptome of stem cell and neural genes. ZFP207 regulates the expression of the spliceosome, and silencing of *Zfp207* leads to aberrant AS patterns. We further describe ZFP207 as a novel RNA-binding protein (RBP), which might directly affect RNA fate. Taken together, this study uncovers the versatile species-specific roles of ZFP207 and the link to co- and posttranscriptional pathways that impact cell-fate decisions of mouse ESCs.

## Results

### Silencing of *Zfp207* impairs proliferation in mouse ESCs

To explore the function of ZFP207 in mouse ESCs, we analyzed the expression of *Zfp207* in retinoic acid (RA)-induced differentiation towards the neural lineage (Fig. 1A) and in spontaneous differentiation of ESCs into the three germ layers by embryoid body (EB) formation (Fig. 1B). Expression of the pluripotency factor *Oct4* (also known as *Pou5f1*) was used to monitor proper cell differentiation. Quantitative PCR with reverse transcription (RT-qPCR) revealed that *Zfp207* was significantly enriched in ESCs compared to differentiated cells (Fig. 1, A and B), and its expression levels gradually decreased along the course of differentiation, correlating with the decrease of ZFP207 and OCT4 protein levels (Fig. 1, C and D).

**Figure 1.**
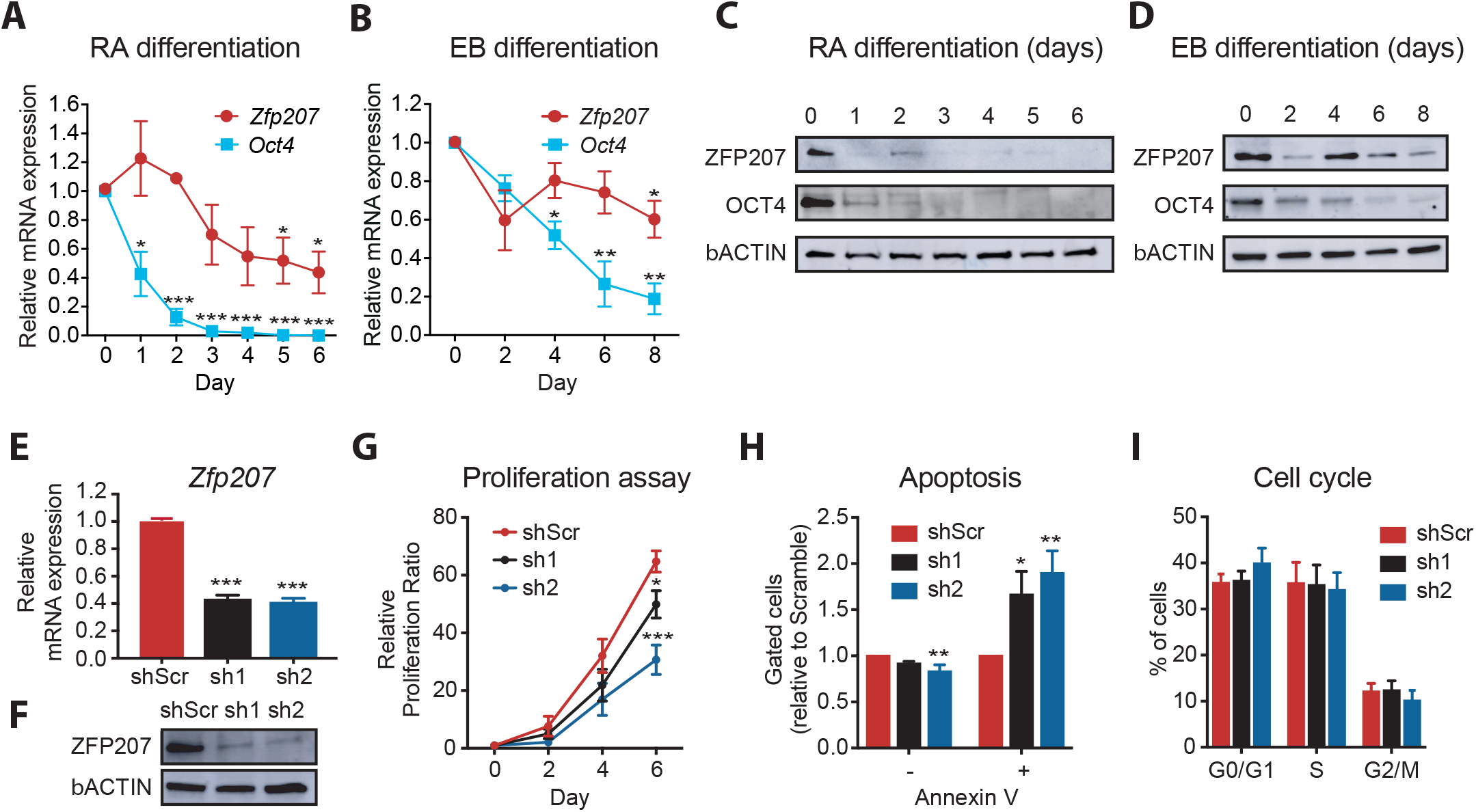
Depletion of *Zfp207* leads to growth defects of mouse ESCs. (**A**) RT-qPCR analysis of *Zfp207* and *Oct4* in mouse ESCs during retinoic acid (RA)-induced and (**B**) embryoid body (EB)-mediated differentiation. mRNA levels are normalized to *β-Actin* and represented as mean ± SEM relative to the expression at day 0. n = 3, * p < 0.05, ** p < 0.001, *** p < 0.0001. An unpaired Student’s *t*-test was used for statistical analysis. (**C**) Representative western blot of ZFP207 and OCT4 of mouse ESCs during RA-induced and **(D)** EB-mediated differentiation. β-ACTIN was used as a loading control. (**E**) RT-qPCR and western blot (**F**) to monitor the knockdown efficiency of *Zfp207* (sh1 and sh2). *Zfp207* is normalized to *β-Actin* and represented as mean ± SEM relative to the expression of the scramble control (shScr). n = 3, *** p < 0.0001. Statistical significance was determined using the ordinary one-way ANOVA. For the immunoblot, β-ACTIN was used as a loading control. (**G**) Cell proliferation rate of shScr, sh1 and sh2 ESCs assessed over a period of 6 days. Measurements were taken every other day. The cell proliferation rates are represented as mean ± SEM, n = 3, * p < 0.05, *** p < 0.0001. Statistical analysis was performed with an unpaired Student’s *t*-test. (**H**) Percentage of live (Annexin V-) and apoptotic cells (Annexin V+) in sh1 and sh2 mouse ESCs compared to shScr. Data are represented as mean ± SEM, n = 3, * p < 0.05, ** p < 0.01. An unpaired Student’s *t*-test was used for statistical analysis. (**I**) Cell cycle distribution in sh1 and sh2 mouse ESCs relative to shScr was examined according to the DNA content index. Data are represented as mean ± SEM, n = 3

To better understand the role of ZFP207 in pluripotency and differentiation, we aimed to generate CRISPR/Cas9-mediated knockout (KO) of *Zfp207* in ESCs. We used two different strategies: (i) one sgRNA (sgRNAs #1) targeting exon 3; and (ii) two distinct single guide RNAs (#1 and #2) targeting the region containing exon 3 and exon 9 which included the microtubule-binding region domain of ZFP207 (fig. S1A). After picking and expanding individual clones, Western blot analysis and subsequent Sanger sequencing confirmed that all clones analyzed (94) displayed heterozygosity for *Zfp207* (fig. S1, B-E) suggesting that homozygous deletion of ZFP207 is lethal as previously suggested (Blomen et al., 2015). Next, we conducted loss-of-function assays by using two distinct short-hairpin RNAs (shRNAs) against *Zfp207* (thereafter referred as knockdown 1 and 2 (KD1 and KD2)), to ensure that the observed phenotype is due to *Zfp207* depletion and not due to shRNA off-target effects. Silencing of *Zfp207* gene expression in both KD1 and KD2 ESCs was confirmed by RT-qPCR (Fig. 1E) and by Western blot (Fig. 1F). Depletion of *Zfp207* led to a reduced proliferation capacity compared to ESCs transduced with scrambled shRNA control (thereafter referred as control), although to a lesser degree in KD1 than in KD2 ESCs (Fig. 1G; 1.3 and 2-fold, respectively). In addition, KD1 and KD2 ESCs showed a significant 1.5-fold increased apoptotic rate compared to control ESCs (Fig. 1H) whilst no significant differences in the cell cycle profile were detected between the three cell lines (Fig. 1I). Overall, our results indicate that ZFP207 is required for the proper proliferation of mouse ESCs.

### ZFP207 maintains stem cell identity

*Zfp207*-depleted colonies displayed the typical morphology of differentiating ESCs with flat appearance and undefined colony borders (Fig. 2A). Consistently, we detected reduced metabolic activity in ESC after depletion of *Zfp207* determined by alkaline phosphatase activity assay (Fig. 2A). Specifically, down-regulation of *Zfp207* resulted in a significant increase in the percentage of partially differentiated colonies whereas the percentage of undifferentiated colonies was significantly decreased compared to control ESCs (Fig. 2B). Such increase in the percentage of partially and finally differentiated colonies in KD1 and KD2 ESCs is a consequence of impaired ESC function as immunofluorescence analysis revealed that silencing of *Zfp207* leads to a decrease of the pluripotency surface marker SSEA1 (Fig. 2C).

**Figure 2.**
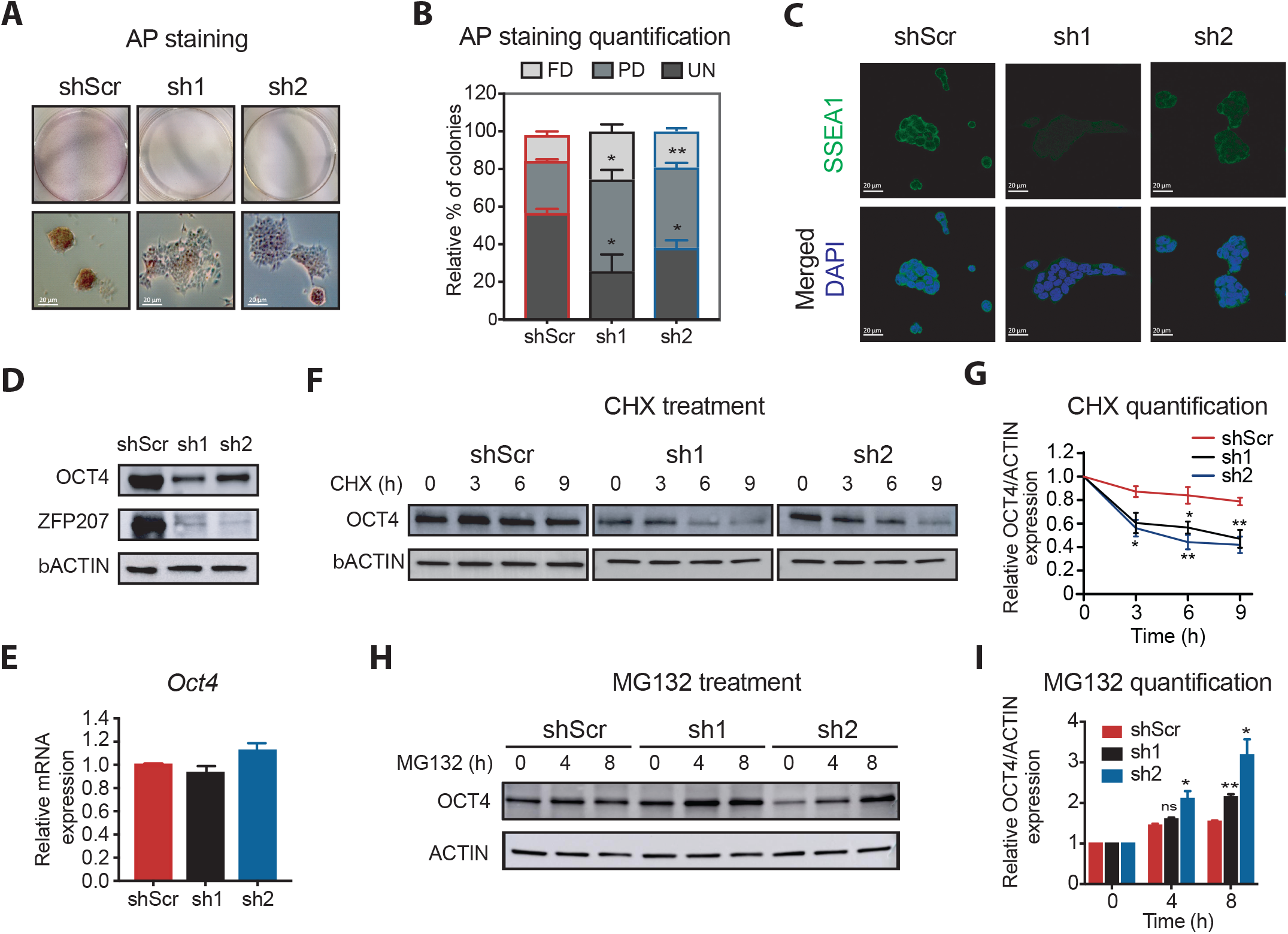
ZFP207 is required for mouse ESCs self-renewal. (**A**) Representative images of AP staining of control (shScr) and *Zfp207-*depleted (sh1 and sh2) mouse ESCs. Scale bars, 20 µM. (**B**) Percentage of fully differentiated (FD), partially differentiated (PD) and undifferentiated (UN) ESC colonies in shScr, sh1 and sh2. Data are represented as mean ± SEM, n = 3, * p < 0.05, ** p < 0.001. An unpaired Student’s *t*-test was used for statistical analysis. (**C**) Representative immunofluorescence analysis of SSEA1 in shScr, sh1 and sh2 ESCs. DAPI was used as the nuclear marker. Scale bars, 20 μm. (**D**) Representative western blot analysis of OCT4 and ZFP207 in shScr, sh1 and sh2 ESCs. β-ACTIN was used as a loading control. (**E**) RT-qPCR analysis of *Oct4* in shScr, sh1 and sh2 ESCs. *Oct4* is normalized to *β-Actin* and represented as mean ± SEM relative to shScr; n = 3. (**F**) Western blot analysis of OCT4 during a 9-h cycloheximide (CHX) time course treatment in shScr, sh1 and sh2 ESCs. β-ACTIN was used as a loading control. (**G**) Protein degradation curves were made after quantification and normalization of the bands from (F). Data are represented as mean ± SEM, n = 3, * p < 0.05, ** p < 0.01. Statistical significance was determined using the ordinary one-way ANOVA. (**H**) Western blot analysis of OCT4 during 4 and 8 h treatment with the proteasome inhibitor MG132. β-ACTIN was used as a loading control and bands were quantified and normalized (**I**). Data are represented as mean ± SEM, n = 3, * p < 0.05, ** p < 0.01, ns = not significative. Statistical significance was determined using the ordinary one-way ANOVA.

Since ZFP207 regulates OCT4 transcription in human, we next analyzed the expression of the pluripotency marker OCT4 and observed that KD1 and KD2 ESCs expressed lower OCT4 protein levels (Fig. 2D) but not *Oct4* mRNA levels (Fig. 2E) compared to control ESCs, indicating that ZFP207 could regulate the expression of OCT4 post-transcriptionally. To test this hypothesis, we treated KD1 and KD2 as well as control ESCs with the protein synthesis inhibitor cycloheximide (CHX). Silencing of *Zfp207* led to a decrease in the half-life of OCT4 (Fig. 2 F and G), suggesting that ZFP207 promotes the stability of this pluripotency factor. In addition, treatment with the proteasome inhibitor MG132 restored OCT4 protein levels in mouse ESCs depleted of *Zfp207* (Fig. 2, H and I), indicating that ZFP207 interferes with the turnover of this pluripotency factor. Altogether, these results suggest that ZFP207 contributes to the maintenance of the pluripotent state in mouse ESC by regulating post-transcriptional networks.

### Silencing of *Zfp207* blocks neuroectodermal differentiation

We next interrogated the role of ZFP207 during lineage specification by assessing the potential of KD1, KD2 and control mouse ESCs to spontaneously differentiated into EBs recapitulating early mouse embryo development. *Zfp207* KD1 and KD2 ESCs were able to form EBs (Fig. 3A and fig. S2A). Although the cells remained as solid aggregates, we observed a decrease in the size of both *Zfp207* KD1 and KD2 EBs compared to control EBs (Fig. 3B), suggesting intrinsic differences during the differentiation process among the different cell lines.

**Figure 3.**
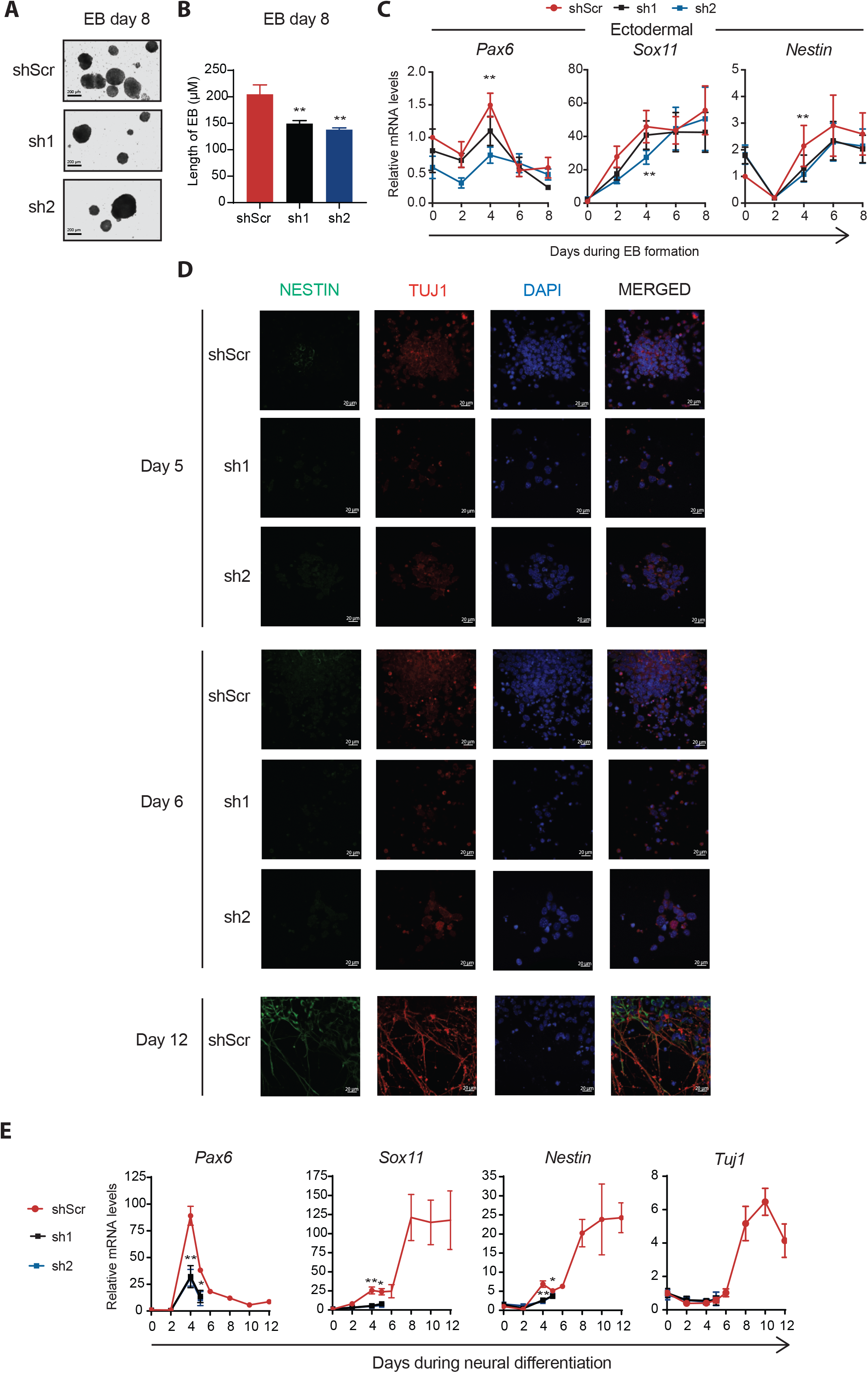
Loss of *Zfp207* results in defective differentiation. Representative bright-field images (10x magnification) of embryoid bodies (EB) generated from control (shScr), and *Zfp207*-depleted ESCs (sh1 and sh2) at day 8 of differentiation. Scale bars, 200 µM. (**B**) Quantification of EB size in sh1 and sh2 ESCs compared to shScr. Data are represented as mean ± SEM, n =3, ** p < 0.001. An unpaired Student’s *t*-test was used for statistical analysis. (**C**) RT-qPCR analysis of the neural-associated genes (*Pax6, Sox11* and *Nestin*) in shScr, sh1 and sh2 ESCs along the time-course of EB-mediated differentiation. mRNA levels are normalized to *β-Actin* and represented as mean ± SEM relative to the expression of shScr at day 0; n =3, ** p < 0.001. A ratio paired Student’s *t*-test was used for statistical analysis. (**D**) Immunostaining of NESTIN (green) and TUJ1 (red) in shScr, sh1 and sh2 derived neural progenitors on the indicated days of neuroectodermal differentiation. Nuclei were counterstained with DAPI. Scale bar, 20 µM. (**E**) Gene expression of the neural-associated markers (*Pax6, Sox11, Nestin, and Tuj1*) in shScr, sh1 and sh2 ESCs along the course of directed neural differentiation. Data points for sh1 and sh2 were profiled up to day 5 because the cells died. mRNA levels are normalized to *β-Actin* and represented as mean ± SEM relative to the expression of shScr at day 0; n =3, * p < 0.05, ** p < 0.001. An unpaired Student’s *t*-test was used for statistical analysis.

As expected, while *Zfp207* expression was gradually down-regulated during the course of differentiation in control cells, *Zfp207* mRNA levels remained low in both KD1 and KD2 cells (fig. S2B). The expression of the pluripotent genes *Nanog* and *Sox2* decreased rapidly in all the cell lines whereas *Oct4* decreased progressively, being its expression higher at the latest stage of the time course differentiation in EBs depleted of *Zfp207* compared to control (fig. S2B). EBs originated from the three-shRNA infected ESCs showed normal levels of the mesodermal (*Msx1* and *Brachyury* or *T*) and endodermal (*Foxa2* and *Sox17*) lineage-specific markers (fig. S2B). However, silencing of *Zfp207* impaired ectodermal specification as shown by decreased expression of *Pax6* and *Sox11* at day four, and *Nestin* during the course of differentiation in *Zfp207* KD EBs compared to controls (Fig. 3C).

To further investigate whether ZFP207 plays a role in neurogenesis, KD1, KD2 and control ESCs were subjected to neural differentiation (fig. S2C). Noticeable, we were not able to characterize *Zfp207* KD1 and KD2 cells at later stages as they died at day 6. Similar to RA- and EB-mediated differentiation, *Zfp207* expression was down-regulated during the course of neurogenesis in control cells, and *Zfp207* mRNA levels remained low in both KD1 and KD2 cells (fig. S2D). The distinct cell lines did not display major morphological differences up to day five of differentiation but KD1 and KD2 underwent a complete block of neuronal differentiation potential at day six (Fig. 3D). In control cell lines, Nestin- and Tuj1-immunopositive neuronal projections appeared at day eight (fig. S2E) and were kept in culture until day twelve (Fig. 3D). We also monitored progression of ESC differentiation toward neural fates by RT-qPCR analysis of neuroectodermal markers. *Zfp207*-depleted cell lines failed to activate the expression of *Pax6, Sox11* and *Nestin* (Fig. 3E and fig. S2F), consistent with a stall at early ectodermal differentiation. No defects were found in the expression of *Tuj1* at the transcriptional level as the up-regulation of this neuronal marker occurred at later stages when the KD1 and KD2 cells had died (Fig. 3E). Altogether these results indicate that ZFP207 expression is required for proper differentiation towards the neuroectoderm lineage.

### ZFP207 does not regulate the ectodermal lineage

In order to gain insight on the role of ZFP207 in ESC pluripotency, we analyzed the global transcriptome response to *Zfp207* depletion (Fig. 4A). RNA-sequencing (RNA-seq) analysis identified 382 and 522 genes that were down-regulated in KD1 and KD2 ESCs, respectively (fold change > 1.5; p < 0.05; Fig. 4B and table S1). The differences of number of down-regulated genes between the two KDs could result in the more severe phenotype observed in cells transduced with shRNA2 compared to shRNA1 (Fig. 1G). The effect on up-regulation was more robust, whereby 1062 and 1082 genes were up-regulated in KD1 and KD2 ESCs, respectively, compared to control ESCs (Fig. 4B). Gene ontology (GO) analysis of biological processes of common down-regulated genes revealed generic functions which included mitotic sister chromatid segregation among other categories (Fig. 4C). Strikingly, poly(A)^+^ RNA binding was amongst the most represented molecular function GO categories (fig. S3A). According to the reported function of ZFP207 in kinetochore-microtubule attachment (Dai et al., 2016), top GO categories for molecular function of common down-regulated genes also included microtubule binding (fig. S3A). We validated these results by performing RT-qPCR analysis of down-regulated mitotic sister chromatid segregation genes (e.g., *Cdca8* and *Cep57l1*; fig. S3B). Up-regulated genes were associated with GO biological processes related to RNA splicing and processing, and positive regulation of transcription (Fig. 4D). Similar to the common down-regulated genes, RNA binding, including poly(A) ^+^ and mRNA binding, were amongst the most represented categories of GO molecular function (fig. S3C), suggesting a role of ZFP207 in controlling the expression of putative RBPs, primarily involved in RNA splicing and mRNA processing (fig. S3D).

**Figure 4.**
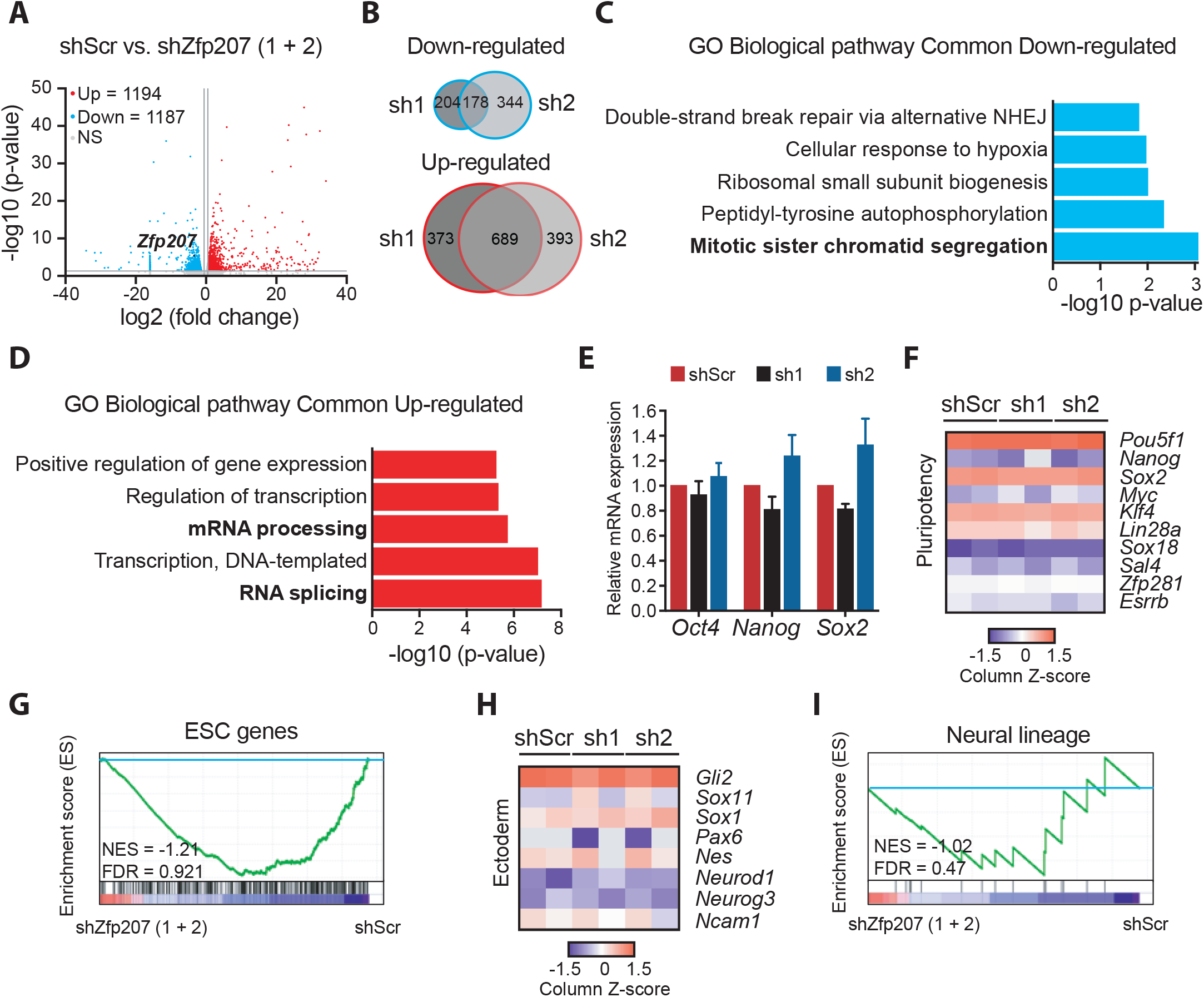
Neural transcriptome is not altered upon depletion of *ZfP207*. (**A**) Volcano plots of common differentially expressed genes in control (shScr) and *Zfp207-* depleted (sh1 and sh2) mouse ESCs. The statistically significant (p < 0.05; > 1.5-fold) up-regulated (Up) and down-regulated (Down) genes are indicated in red and blue, respectively. *Zfp207* is depicted in black. Grey dots indicate non-significant (NS) and < 1.5-fold differential expressed genes. (**B**) Venn diagram depicting the overlap of down-regulated and up-regulated genes between sh1 and sh2 ESCs (FC>1.5 and p < 0.05). (**C**) Gene ontology (GO) analysis of biological processes associated with common down-regulated and (**D**) up-regulated genes in *Zfp207*-depleted ESCs (sh1 and sh2); NHEJ: Non-homologous end joining. (**E**) RT-qPCR analysis of pluripotency markers *(Oct4, Nanog* and *Sox2)* in shScr, sh1 and sh2 ESCs. mRNA expression is normalized to *β-Actin* and represented as mean ± SEM relative to shScr; n = 3. (**F**) Heatmap of column z-scores of log2 transformed values of genes related to pluripotency in shScr, sh1 and sh2 ESCs. (**G**) GSEA plots depicting the expression of ESC genes in *Zfp207-* depleted ESCs compared to shScr. High and low expression of genes is represented in red and blue color, respectively. FDR, false discovery rate; NES, normalized enrichment score. (**H**) Heatmap and (**I**) GSEA plot showing the global expression of genes related to ectoderm and neural lineage in *Zfp207-*depleted ESCs compared to shScr. High and low expression of genes is represented in red and blue color respectively. FDR, false discovery rate; NES, normalized enrichment score.

Depletion of *Zfp207* did not lead to aberrant transcriptional programs that could explain the developmental defects. Hence, the core pluripotency regulators *Oct4, Nanog* and *Sox2*, were not differentially expressed in KD1 and KD2 ESCs compared to control ESCs (Fig. 4E), and there were no differences in the expression of other ESC genes upon silencing of *Zfp207* (Fig. 4, F and G). In addition, we did not find major transcriptional differences in genes associated with the neural lineage that could reflect the stall at early ectodermal differentiation in KD1 and KD2 ESCs (Fig. 4, H and I). This is in striking contrast to what it has been reported in human ESCs where depletion of *ZNF207* impairs neuroectodermal specification by transcriptionally regulating the expression of genes associated with the ectoderm lineage (Fang et al., 2018). Noteworthy, in human ESCs the isoforms A and C of *ZNF207*, retaining the exon 9, are highly abundant compared to mouse ESCs, where the isoforms 1 and 2 of *Zfp207*, retaining the aforementioned exon 9, are highly expressed in differentiated cells (fig. S3, E and F). Hence, in both species there is an antagonistic switch towards using different isoforms during differentiation which could partially explain the differences observed between mouse and human ESCs.

### ZFP207 regulates alternative splicing in mouse ESCs

AS is a fundamental process that increases proteomic diversity in a species-but not organ-specific manner (Barbosa-Morais et al., 2012), and it also plays a central role in the regulation of ESC-specific transcriptional programs (Gabut et al., 2011). ZNF207 has been shown to influence pre-mRNA splicing in cancer cells (Wan et al., 2015). Consistent with this observation, our GO enrichment analysis of genes up-regulated upon *Zfp207* silencing revealed significant enrichment of GO terms related to RNA splicing (Fig. 4D and Fig. 5A), and some randomly selected spliceosomal genes (*Mbnl2, Rbm3* and *Sf3b3*) were further validated by RT-qPCR (Fig. 5B).

**Figure 5.**
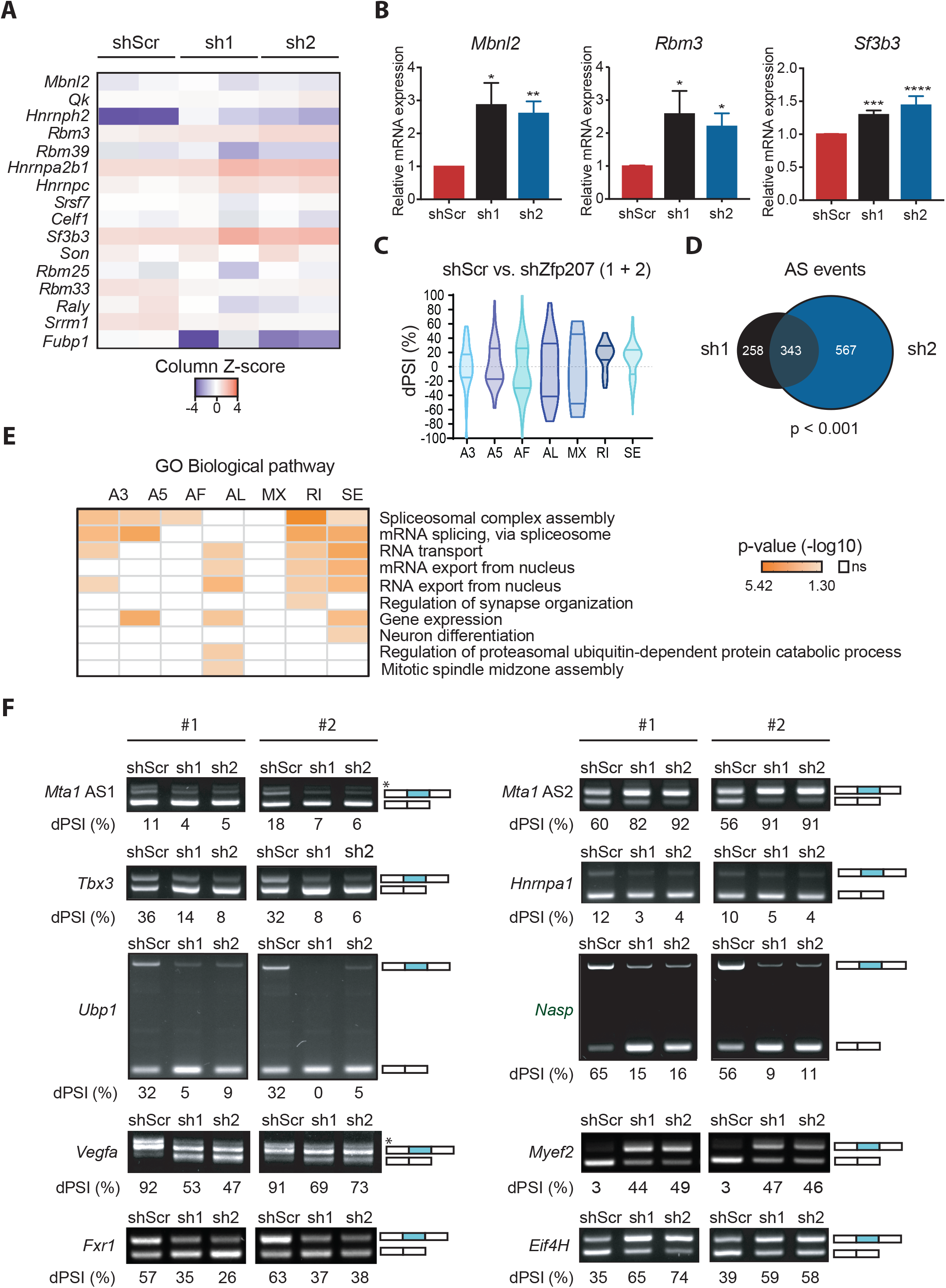
Silencing of *Zfp207* leads to AS defects. (**A**) Heatmap of column z-scores of log2 transformed values of genes involved in alternative splicing in control (shScr) and *Zfp207-*depleted (sh1 and sh2) mouse ESCs. (**B**) RT-qPCR analysis of *Rbm3, Mbnl2* and *Sf3b3* in shScr, sh1 and sh2 ESCs. mRNA expression is normalized to *β-Actin* and represented as mean ± SEM relative to shScr; * p < 0.05, ** p < 0.01, *** p < 0.001, **** p < 0.0001, n = 3. An unpaired Student’s *t*-test was used for statistical analysis. (**C**) Violin plot depicting AS events in each category in mouse ESCs depleted of *Zfp207* (sh1 and sh2) related to shScr. A3 (Alternative 3’); A5 (Alternative 5’); AF (Alternative first exon); AL (Alternative last exon); MX (Mutually exclusive exons); RI (Intron retention); SE (Exon skipping). (**D**) Venn diagram showing the overlap of genes undergoing AS in mouse ESCs infected with sh1 and sh2 against *Zfp207*. (**E**) Gene ontology analysis of common genes undergoing AS in mouse ESCs infected with sh1 and sh2 against *Zfp207* according to the type of AS event. (**F**) Representative RT-PCR validations of *Zfp207*-regulated AS events (*Mta1*; *Tbx3*; *Ubp1*; *Vegfa*, and *Fxr1* (left panel) and Mta1, *Hnrnpa1, Nasp, Myef2, Eif4H* (right panel)). For *Mta1*, AS1 (alternative splicing 1) and AS2 (alternative splicing 2). #1 and #2 indicate distinct biological replicates. *Denotes an isoform which was not taken in consideration for the quantification. The structure of each isoform is indicated (not to scale). Alternative exons are blue. The percent spliced in (PSI) was quantified for each condition.

We therefore sought to investigate whether ZFP207 could control ESC function by a splicing-related mechanism. To this end, reads from RNA-seq data were mapped to exon-splice junction’s sites in order to elucidate genome-wide differential AS events (DSEs) (Alamancos et al., 2015; Trincado et al., 2018b), including alternative 5’ and 3’ splice-site selection (A5 or A3), alternative first (AF) and last (AL) exon selection, exons that are spliced in a mutually exclusive manner (MX) or that are skipped (SE), and changes in intron retention (RI) (fig. S4A). Comparisons of AS isoform levels were performed in KD1 and KD2 ESCs versus control shRNA ESCs, and differences in AS isoforms were calculated as the change in percent spliced in (dPSI) (Fig. 5C). Alternative first exon selection and exon skipping appeared to be the most predominant splice events (fig. S4B). Our analysis identified 601 and 910 splicing events in KD1 and KD2 compared to control corresponding to 419 and 609 genes, respectively (false discovery rate [FDR] < 0.05; Fig. 5D and table S2). Alterations in DSEs upon knockdown of *Zfp207* were not due to changes in transcription as gene expression levels were not correlated to dPSI (fig. S4C). The overlapped AS genes were significantly enriched in functional categories associated with RNA splicing, RNA transport and export from the nucleus, and neurogenesis, among others (Fig. 5E and fig. S4D), suggesting that splicing switches occurring upon silencing of *Zfp207* are relevant to the severe blocking defect towards the neural lineage observed in KD1 and KD2 ESCs. DSEs were validated by RT-PCR amplifying AS exons that differ in size and band intensity was assessed in order to estimate the exon-inclusion ratios. Specifically, we validated randomly selected ZFP207-mediated DSEs in control, KD1 and KD2 ESCs (Fig. 5F), including the chromatin regulatory cofactor *Mta1* (AS1 and AS2); the AS factors *Tbx3* and *Hnrnpa1*; the transcription factor *Ubp1*; the histone chaperone *Nasp*; the neuronal factors *Vegfa, Myef2* and *Fxr1*; and the translation initiation factor *eIF4H*. Furthermore, we also interrogated the aforementioned DSEs in neural differentiation, and found that the majority of the splicing switches validated in KD1 and KD2 ESCs were also prevalent in differentiated cells (fig. S4E). Collectively, these data indicate that depletion of *Zfp207* elicits aberrant AS patterns that resemble the differentiated cell-like pattern, and consequently influences mouse ESC identity.

### ZFP207 is a newly identified RBP

Because ZFP207 influenced AS and it has been identified as a mRNA binder using a quantitative proteomics approach (Baltz et al., 2012; Castello et al., 2012), we assessed the innate binding propensity of ZFP207 in mouse ESCs by employing a modified *in vitro* RIP protocol (see methods section). We identified 1,400 transcriptome-wide binding events for ZFP207 in our assay (table S3). In total, 942 of these binding sites resided within coding regions of protein-coding genes (Fig. 6A). Since ZFP207 is located in the nucleus, we also found binding to intronic sequences (178/1,400). In addition, we detected binding to untranslated regions (UTRs) (267/1,400) and exons of non-coding transcripts (17/1,400). It is possible that there are more ZFP207-binding events to nascent RNAs, which might have been missed since most intronic sequences are depleted by this method. The 1,400 ZFP207 binding events were distributed over 1,171 genes. Based on a fold-change weighted by the relative expression levels (referred to as binding score), we identified the strongest binding sites within the *Ubp1, Marf1, Sae1, Erp29* and *Nsd1* transcripts (Fig. 6B). Surprisingly, three of the top five binding sites (*Marf1, Sae1* and *Nsd1*) were found over exon-exon junctions (fig. S5A). Such binding events are difficult to envision *in vivo*, unless ZFP207 is part of the exon-exon junction complex and connects exonic ends directly.

**Figure 6.**
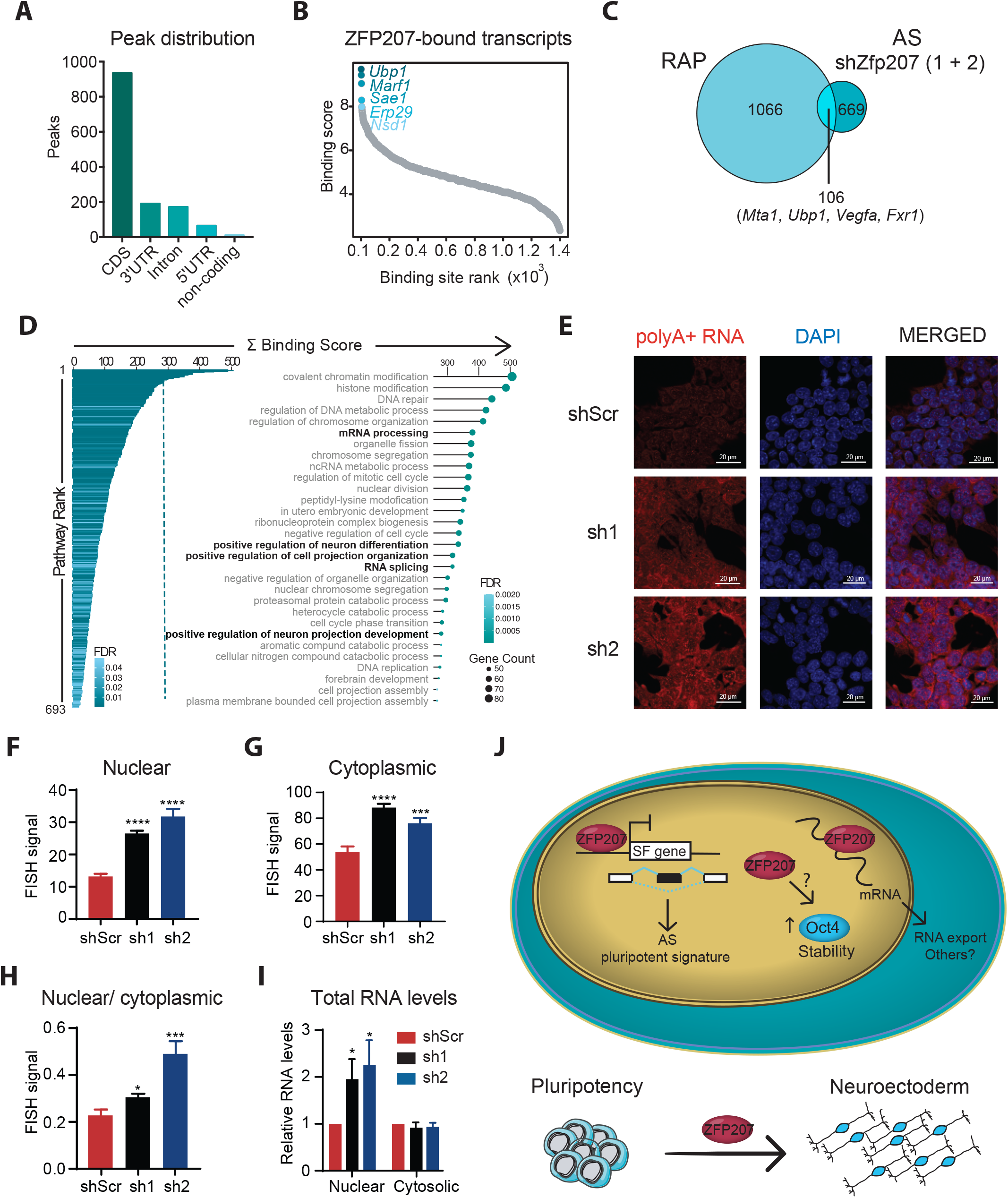
ZFP207 is a novel RBP. (**A**) Schematic representation of the RNA Affinity Purification followed by Sequencing (RAPseq) experimental procedure. (**B**) Barchart representing the distribution of ZFP207 binding sites across common transcript features. (**C**) Scatterplot depicting ranked distribution of ZFP207 binding sites. The top 5 strongest binding sites are highlighted. (**D**) Venn diagram showing the overlap of genes retrieved by RAP-seq and genes undergoing AS in mouse ESCs upon *Zfp207* knockdown. (**E**) Bar- and dot plots show significantly enriched GO terms for Biological processes for ZFP207 bound genes. Each term is ranked by the sum of all binding scores of all peaks present in this pathway. Top 30 GO terms are shown. Circle diameter represents the number of genes bound by ZFP207 in that specific pathway. **(F)** FISH analysis of poly(A)^+^ RNA distribution in scramble control (shScr) and *Zfp207-*depleted ESCs (sh1 and sh2) stained with 5’Cys3-oligo dT(50). Nuclei were counterstained with DAPI. All images were subjected to the same exposure settings. Scale bar, 20 µM. **(G, H)** Quantification of nuclear and cytoplasmic poly(A)^+^ RNA FISH signals in shScr, sh1 and sh2 respectively. Data are represented as mean ± SEM; *** p < 0.001, **** p < 0.0001; n = 2. An unpaired Student’s *t*-test was used for statistical analysis. **(I)** Ratio of nuclear/cytoplasmic poly(A)^+^ FISH signals in shScr, sh1 and sh2 mouse ESCs. Data are represented as mean ± SEM; * p < 0.05, *** p < 0.001; n = 2. An unpaired Student’s *t*-test was used for statistical analysis. **(J)** Relative nuclear and cytosolic total RNA levels in *Zfp207-*depleted ESCs compared to shScr. Results are represented as mean ± SEM; * p < 0.05; n = 2. An unpaired Student’s *t*-test was used for statistical analysis. **(K)** ZFP207 modulates mouse ESCs pluripotency and neuroectodermal differentiation. In our model, ZFP207 regulates the expression of multiple splicing factors and AS pattern, influencing cellular fate specification. Besides, ZFP207 regulates OCT4 stability post-transcriptional networks. In addition, ZFP207 acts as an RBP to promote mRNA nuclear export. Hence, the co- and post-transcriptional regulatory mechanisms controlled by ZFP207 are essential for maintaining proper mouse ESC identity.

Overlapping of the RAPseq and AS data sets revealed 13.7% of DSEs that were directly bound by ZFP207 (Fig. 6C), including *Mta1, Ubp1, Vegfa*, and *Fxr1* with AS switches occurring upon silencing of *Zfp207* that were validated (Fig. 5F). However, we cannot exclude that dysregulated expression of genes encoding spliceosomal factors upon depletion of *Zfp207* was a major contributor to ESC-differential AS events.

In order to classify the bound genes into functional groups, we performed a pathway enrichment analysis for biological processes and we found that the 1,171 genes felt into 693 significantly enriched pathways over the whole murine transcriptome (FDR ≤ 0.05) (Fig. 6D). Interestingly, GO analysis revealed DNA repair, chromosome organization and chromosome segregation among the most represented categories, suggesting that ZFP207 can also regulate the aforementioned pathways not just by interacting with Bub3 (Jiang et al., 2014; Toledo et al., 2014) but also by directly binding to target transcripts and influencing RNA fate. Noteworthy, GO terms that were affected upon *Zfp207* silencing such as mRNA processing, splicing, positive regulation of neuron differentiation, and neuron projection development, were highly enriched. This suggests that in addition to regulating a pluripotent AS signature, ZFP207 controls the ESC state through post-transcriptional gene regulatory mechanisms. Indeed, ZFP207-bound transcripts felt into two major ESC gene signatures, namely the Core and Myc modules, which are highly active in the pluripotent state, whereas the Polycomb repressive (PRC) module, operating in differentiation, was under-represented (fig. S5B) (Kim et al., 2010), again in agreement with the observation that ZFP207 regulates pluripotency by transcriptional-independent programs. The fact that ZFP207-bound transcripts were enriched in the Myc module, which is active in various cancers and predicts cancer outcome, may reveal additional post-transcriptional regulatory functions of ZFP207 not just in pluripotency but also in human diseases. Depiction of the binding score of the ZFP207-bound transcripts showed similar binding activity pattern within the three modules (fig. S5C).

Because RBPs play important roles not just in mRNA splicing but also in other aspects of the posttranscriptional mRNA fate such as nuclear export, localization or nuclear decay (Ye and Blelloch, 2014), we analyzed the distribution of bulk poly(A)^+^ RNAs by Fluorescence in situ hybridization (FISH) using an oligo(dT) probe in scramble control, KD1 and KD2 ESCs. mRNA was accumulated in nuclear poly(A)^+^ foci upon silencing of *Zfp207* (Fig. 6, E and F), suggesting that ZFP207 might play a role in nuclear export. Although cytoplasmic poly(A)^+^ RNA FISH signals were also brighter in *Zfp207-*depleted cells compared to that in control cells (Fig. 6, E and G), the intensity of cytoplasmic poly(A)^+^ RNA pixel and the nuclear to cytoplasmic ratios were less profound in *Zfp207* knockdown compared to control ESCs (Fig. 6, G and H). In addition, cellular fractionation showed a clear accumulation of nuclear RNA upon silencing of *Zfp207* whereas no differences were detected in the amount of RNA in the cytoplasmic fractions of the distinct cell lines (Fig. 6I). Although it remains possible that some mRNA will get exported from the nucleus, these results indicate that at least some particular mRNAs are nuclear retained and largely aggregate into poly(A)^+^ foci in *Zfp207* KD1 and KD2 ESCs.

In summary, we show that ZFP207 plays an important role in maintaining mouse ESC self-renewal and neuroectodermal specification by controlling co- and post-transcriptional networks. We propose a model whereby ZFP207 regulates the expression of splicing factors to promote *bona fide* splicing of a specific set of transcripts that alter cellular fate. Additionally, ZFP207 acts as an RBP to facilitate mRNA nuclear export. Hence, the versatility in regulatory capacities of ZFP207 are important for normal expression of pluripotency and neural commitment genes in order to ensure mouse ESC identity (Fig. 6J).

## Discussion

Previous investigations have focused attention on the properties of ZNF207 as a transcription factor and its function in transcriptional control (Fang et al., 2018). An important aspect of our findings is the observation that ZFP207 regulates mouse ESC maintenance through co- and post-transcriptional mechanisms. On the contrary to as human ESCs, mouse ESCs have a naïve pluripotency with unbiased differentiation capacity, and they have proven to be an invaluable model to study development. Hence, although the human orthologue ZNF207 has been described as a critical regulator for self-renewal and pluripotency elsewhere (Fang et al., 2018), we found divergent functional characteristics between both studies. These dissimilarities may reflect two distinct pluripotent states that represent mouse and human ESCs: naïve and primed, respectively (Ginis et al., 2004). In addition, these differences could arise from opposite AS shifting patterns of *Zfp207* and *ZNF207* that occur during cellular differentiation of mouse and human ESCs, respectively.

ZFP207 is essential for cellular viability in near-haploid cells (Blomen et al., 2015). In line with this, we were unable to fully disrupt the *Zfp207* gene in mouse ESCs using CRISPR/Cas9 technology. Hence, we anticipate that ablation of ZFP207 is incompatible with ESC maintenance. Moreover, our results show that depletion of *Zfp207* in ESCs resulted in cell proliferation defects by triggering apoptosis, which further shifts the self-renewal phenotype towards differentiation. Similar to human ESCs, we also found that ZFP207 controlled the expression of the pluripotency factor OCT4. However, such regulation did not occur transcriptionally as *Oct4* mRNA levels were not affected upon silencing of *Zfp207*. OCT4 is target for ubiquitination and degradation, and this post-translational modification plays a critical role in OCT4 expression in ESCs (Xu et al., 2009). Our data suggests that ZFP207 would increase OCT4 protein stability by preventing degradation of OCT4 through the proteasome. Consistently, Jiang et al. (Jiang et al., 2014), reported that ZFP207 functions as a chaperone that binds and stabilizes the spindle assembly checkpoint protein Bub3 by protecting Bub3 from degradation by the proteasome. It remains to be elucidated the mechanism by which ZFP207 protects OCT4 from proteasomal degradation as none of the core pluripotency factors were identified in mass spectrometric analysis of mouse ESCs using ZFP207 as bait (Jiang et al., 2014). However, such study identified the E3 ubiquitin ligase UBR5 as a ZFP207-interacting partner, suggesting that ZFP207 could exert a secondary role in controlling proteasomal-mediated degradation.

We assessed the function of ZFP207 during *in vitro* differentiation to the three germ layers, recapitulating the early events in embryogenesis, by performing EB-directed differentiation. Although EB formation and growth were defective after silencing of *Zfp207*, we could conclude from these experiments that endodermal and mesoderm specification were normal whereas the neuroectodermal lineage was highly compromised. Indeed, upon neural induction, EBs depleted of *Zfp207* failed to efficiently up-regulate neural markers, and neuronal differentiation appeared to stall at an early stage. In contrast to human ESCs, RNA-seq analysis revealed that ZFP207 did not drive the transcription of neuronal gene expression programs and could not explain the blockade of neuroectodermal differentiation upon *Zfp207* silencing. However, ZFP207 controlled the transcription of multiple splicing factors which potentially could lead to aberrant AS patterns with critical functions in pluripotency. For instance, we validated the up-regulated expression of *Mbnl2, Rbm3, and Sf3b3*. Accordingly, *Mbnl2* and *Rbm3* have been shown to promote differentiated-cell-specific AS patterns (Han et al., 2013; Xia et al., 2018; Yan et al., 2019). To our knowledge, no specific function of the splicing factor Sf3b3 in ESCs has been described. Global profiling of AS landscape revealed alterations of a broad number of genes. Although such splicing defects could be consequence of an indirect effect, through dysregulation of spliceosome gene expression, we cannot exclude the possibility that ZFP207 also regulates AS in a direct manner, through interaction with spliceosome factors (Jiang et al., 2014). Hence, it has been shown that ZFP207 interacts with the splicing components U2AF65 and SF3a3, and transcriptomic analysis revealed splicing defects after depletion of *ZFP207* in cancer cells, although such splicing events were not thoroughly characterized (Wan et al., 2015). Our results expand on these prior observations reporting that ZFP207 is a major splicing regulator that plays a critical role in modulating post-transcriptional networks that influence cell fate.

Functional categories of these AS transcripts included RNA metabolism and neuron differentiation. Interestingly, mitotic spindle midzone assembly was also amongst the top identified GO categories, suggesting that ZNF207 might not only promote spindle assembly by undergoing coacervation and hence, concentrating microtubules and tubulin through ZNF207 droplets (Jiang et al., 2015), but also by an RNA-mediated mechanism. Randomly chosen differentially spliced events have been experimentally validated in this study. Most of them (eight out of ten), displayed a switching to a differentiated cell-like AS pattern upon *Zfp207* knockdown, indicating that ZFP207 regulates an AS signature that controls pluripotency. In addition, several of the stem cell switches that we identified, such as *Nasp* or *Tbx3*, have been shown to be involved in cancer (Alekseev et al., 2011; Krstic et al., 2020).

By using a modified *in vitro* RIP protocol, we characterized ZFP207 as a novel RBP. ZFP207 has an annotated C2H2-ZNFs through which RNA binding is facilitated (Brannan et al., 2016),and two independent interactome capture studies have retrieved ZFP207 in the mRNA-bound proteome (Baltz et al., 2012; Castello et al., 2012). As the extent of intronic binding events that ZFP207 could potentially have inside the nucleus can be orders of magnitude higher than the ones we identified with the *in vitro* RIP, it would therefore be tempting to speculate that ZFP207 regulates AS by directly binding to specific transcripts. However, it is also plausible that the RNA-binding activity of ZFP207 provide an additional function as we observed an accumulation of poly(A)^+^ foci in KD1 and KD2 ESCs. Interestingly, it has been shown that the poly(A)^+^ RNA foci that were accumulated upon depletion of *Zfp207* are primarily formed by exosome target mRNAs (Fan et al., 2018). Thus, it is possible that ZFP207 functions in the turnover of poly(A)^+^ RNA in general, and in the export of pluripotency and neuroectodermal mRNAs in ESCs. It would be interesting to determine in future studies whether the distinct *Zfp207* isoforms target a distinct set of RNAs. If that would be the case, the ESC-specific switch in the exon 9 could drive divergent RNA-metabolism-based programs required for ESC self-renewal and pluripotency.

Collectively, this study further expands the function of ZFP207 and contributes to our understanding of how co- and posttranscriptional programs ensure gene expression to maintain the ESC state. Given that ZFP207 binds both DNA and RNA, the relative contributions between transcriptional and posttranscriptional networks cannot be discriminated. Hence, we cannot exclude the possibility that ZFP207 may also regulate self-renewal transcriptionally i.e., by regulating the expression of splicing factors. Thus, ZFP207 provides an extra layer of control to fine tune the balance between self-renewal and neuronal lineage commitment in mouse ESCs.

## Materials and Methods

### Antibodies

The following commercially available antibodies were used at the indicated concentrations for western blot: Anti-ZFP207 (Santa Cruz Biotechnology, sc-271943, 1:500), anti–β-ACTIN (Sigma-Aldrich, A5441, 1:2500), Anti-LAMIN A/C (Abcam ab108922), Anti-OCT3/4 (Santa Cruz Biotechnology, sc8628, 1:2500), Goat Anti-Rabbit IgG H&L (HRP) (Abcam, ab6721, 1:5000), Goat Anti-Mouse IgG H&L (HRP) (Abcam, 1:1000, ab6789), Rabbit Anti-Goat IgG H&L (HRP) (Abcam, 1:5000, ab6741), and Rabbit Anti-goat IgG (HRP) (Abcam, ab6771, 1:5000). For IF staining, we used Anti-SSEA1 (ThermoFisher Scientific, MA5-17042, 1:250), Anti-Nestin (Abcam, ab6142, 1:50), Anti-Tuj1 (Abcam, ab18207, 1:200), Anti-OCT3/4 (Santa Cruz Biotechnology, sc8628, 1:2500), Nucleolin (Abcam, ab50279, 1:130), Goat anti-mouse IgG AF488 (ThermoFisher Scientific, A11029, 1:1000), Goat anti-rabbit IgG AF568 (ThermoFisher Scientific, A11011, 1:1000), Anti-Mouse IgG HRP (Abcam, ab6789, 1:1000), and Donkey Anti-Goat IgG AF594 (ThermoFisher Scientific, A11058, 1:250).

### Cell culture

CCE murine ESCs were cultured on 0.1% gelatin-coated tissue culture plates under feeder free culture conditions at 37°C with 5% CO^2^ in a humidified incubator. The media composition consists of Dulbecco’s modified Eagle’s medium (DMEM) high glucose, 15% fetal bovine serum (FBS; Gibco), 1% MEM non-essential amino acids (Sigma-Aldrich), 0.1 mM 2-β mercaptoethanol, 1% L-glutamine (Hyclone) and 1% penicillin/streptomycin (Gibco) and Leukemia inhibitory factor (LIF; R&D systems).

### Lentiviruses production and generation of *Zfp207* KD mouse ESCs

Lentiviral particles were generated by transfecting HEK-293T with the following lentiviral plasmids: i) pLKO.1-Puro containing shRNA1 and shRNA2 against *Zfp207* (table S4); ii) the packaging vector pCMV-dR8.2; and iii) the helper plasmid pCMV-VSV-G using jet-PEI (Polyplus) as per the manufacturer’s instructions. Lentiviral supernatants were collected after 48 hours of incubation and concentrated using Amicon Ultra-15 Centrifugal Filter Units (Merck). Early passage mouse ESCs were transduced with the lentiviral particles in mouse ESC media supplemented with polybrene (8 µg/ml) for 24 hours. After 36 hours, infected cells were treated with puromycin (2 µg/ml) for additional 6 days.

### Generation of CRISPR/Cas9 knockout ESCs

sgRNAs were designed using E-CRISP online tool (http://www.e-crisp.org/E-CRISP/aboutpage.html) (table S4) (Ran et al., 2013). All sgRNA-Cas9 plasmids were obtained by ligation (T7 DNA ligase, Fermentas) of annealed complementary oligonucleotides of the 20-nucleotides target sequences with the pSpCas9(BB)-2A-Puro (PX459) vector (Addgene plasmid #62988) digested with BbsI (BpilI) (Thermo Scientific). One day before transfection, mouse ESCs were seeded in a 12 wells-plate at a density of 80,000 cells/well. The day after, cells were transfected using Lipofectamine (Invitrogen) with 0.8 µg of Cas9 expression vector containing the corresponding sgRNAs. After 24 hours, transfected cells were diluted and treated with puromycin to obtain isogenic cell clones. Isogenic cell clones were picked and expanded for 10 to 15 days to identify indels (insertion and deletion). KO were screened with PCR, Sanger sequencing and evaluated by western blot analysis.

### EB and RA differentiation assays

EBs were obtained by growing mouse ESCs in low-attachment dishes in the presence of complete medium (DMEM high glucose, 15% FBS (Gibco), 1% MEM non-essential amino acids (Sigma-Aldrich), 0.1 mM 2-β mercaptoethanol, 1% L-glutamine (Hyclone) and 1% penicillin/streptomycin (Gibco) without LIF at a density of 8.8 ×10^4^ cells/cm^2^. Medium was replenished every 48 hours. EBs were harvested for extraction of total RNA, whole cell extract and nuclear protein extraction at the indicated time points. RA differentiation was performed plating mouse ESCs onto tissue culture dishes pre-coated with 0.1% gelatin at a density of 2.1 ×10^3^ cells/cm^2^ in complete media supplemented with 1 μM RA (Sigma-Aldrich) for 6 days. Differentiation media containing RA was replenished every 48 hours.

### Neural differentiation assay

Differentiation towards the neuroectodermal lineage was performed as previously described (Aguilo et al., 2017). Briefly, 5×10^4^ cells/ml were plated in low-attachment dishes in the presence of complete medium without LIF. Following EB formation for 2 days, EBs were cultured in media containing DMEM/F-12, 0.1 mM 2-mercaptoethanol, N2 supplement (100X) (ThermoFisher Scientific), B27 supplement (50X) (ThermoFisher Scientific), and penicillin/streptomycin (ThermoFisher Scientific) supplemented with 1 μM RA (Sigma-Aldrich) for 4 days to allow neuroectodermal differentiation. On day 6, EBs were transferred onto tissue culture dishes pre-coated with 0.1% gelatin and neuroectodermal differentiation media was replenished every 48 hours. Differentiated cells were harvested for total RNA, nuclear protein extraction and immunostaining analysis at the indicated time points. Cell cultures were maintained at 37°C with 5% CO^2^ in a humidified incubator.

### Cellular proliferation, cell cycle and apoptosis assays

Cellular proliferation, cell cycle and apoptosis assays were carried out using a Muse Cell Analyzer (Millipore, Sigma-Aldrich) following the manufacturer’s recommendations.

### Alkaline Phosphatase Staining

Alkaline Phosphatase Staining was measured using the Alkaline Phosphatase Detection kit (Stemgent) according to standard protocols from the manufacturer. Brightfield images were obtained using a Zeiss microscope and analysed by ZEN lite software.

### Immunocytochemistry analysis

Cells were fixed in 4% formaldehyde (Sigma-Aldrich) and permeabilized with 0.25% Triton-X-100 (Sigma-Aldrich). Following permeabilization, cells were washed twice with PBS and blocked with 10% goat serum (Invitrogen) and bovine serum albumin (Hyclone) at RT for 1 hour. Cells were then stained with primary antibodies overnight at 4°Cat corresponding dilution as previously indicated and secondary staining was performed using corresponding secondary antibodies. 4’, 6-diamino-2-phenylindole (DAPI) was used for DNA nuclear stain. Images were acquired using a Nikon microscope (ZEN lite software) and analysed using ImageJ.

### CHX treatment and Proteasome inhibition

Mouse ESCs were seeded at a density of 1×10^5^ cells/ml in 6-well plate and after 24 hours, cells were treated with cycloheximide (CHX) at a final concentration of 50 μg/ml for 0, 3, 6, and 9 hours, respectively. For proteasome inhibition, cells were incubated with MG-132 at a final concentration of 10 μg/ml for 0, 4 and 8 hours, respectively. Cells were harvested for whole cell extracts at indicated time points and subjected to immunoblotting. Blots were quantified using Image Lab software.

### RT-qPCR

Extraction of total RNA was performed using RNeasy Plus Mini Kit (Qiagen). One μg of total RNA was used for complementary DNA synthesis using RevertAid First Strand cDNA Synthesis kit (Invitrogen) as per the manufacturer’s instructions. RT-qPCR was performed in triplicate using the PowerUp SYBR Green Mix (ThermoFisher Scientific) and the primers listed in table S4 on a Quantastudio Real-time PCR instrument (Applied Biosystems). ☐-*actin* levels were used to normalize input amounts. After each round of amplification, melting curve analysis was carried out in order to confirm the PCR product specificity. Ct values obtained for each gene were analyzed using the ΔΔCt method from at least three biological replicates.

### RNA-Seq and differential gene expression analysis

RNAseq library preparation was carried out at Novogene facilities (https://en.novogene.com/) and sequenced using Illumina HiSeq 2500 platform (Illumina) as 150 bp pair-ended reads. FASTQ reads were aligned to ENSEMBL mouse transcriptome using BWA-MEM (Li, 2013). Then, the calculation of normalized reads per transcript was retrieved by Express (Roberts and Pachter, 2013). Thereafter, DEG were obtained by using a test under the assumption of a negative binomial distribution where the variance is linked to the mean via a locally-regressed smooth function of the mean (Anders and Huber, 2010) and p-values were adjusted by estimation of the false discovery rate for multiple hypothesis (Benjamini and Hochberg, 1995). We only considered the genes with TPM>0 in at least 2 of the samples. FDR value was calculated with the Benjamini-Hochberg correction.

### AS and data analysis

SUPPA2 was used as pipeline for AS analysis (Patro et al., 2017). Briefly, FASTQ reads from RNAseq experiment were aligned and pseudo quantified to mouse genome (mm10) using Salmon (Trincado et al., 2018a). Splicing events in the mouse genome were obtained using a specific SUPPA2 script from a mouse GTF file. Thereafter, percent splicing inclusion (PSI) values for each event were obtained, and the differential PSI values for each condition was calculated along with a p-value for each event.

### RT-PCR

Total RNA was extracted using the RNeasy Mini Kit (Qiagen). Two μg of total RNA was reverse transcribed using the RevertAid First Strand cDNA Synthesis kit (Invitrogen). RT-PCR was performed using the DreamTaq Green Master mix (Thermo Fisher Scientific). The specific primers used for AS are listed in table S4. The ratios between splice variants were determined by densitometry using ImageJ software. The percentage of inclusion was calculated by dividing the area of the inclusion band by the combined area of both inclusion and exclusions bands and multiply by 100.

### Gene Ontology (GO) Analysis

Gene ontology (GO) analysis was performed using the web tool The Database for Annotation, Visualization and Integrated Discovery (DAVID) (http://david.abcc.ncifcrf.gov/).

### Heatmaps

Heat maps were made using Z-score which was calculated as following: Z=(X-Xav)/Xsd, where X is the log_2_-transformed expression level for a given gene in a specific sample, Xav is the mean of log_2_-transformed expression values for that gene in all samples, and Xsd is the standard deviation of log_2_-transformed expression values for that gene across all samples.

### *In vitro* RIP and data analysis

cDNA of *Zfp207* isoform 3 from mouse ESCs was generated (RevertAid® First Strand cDNA Synthesis Kit) and cloned into a HaloTag backbone as a ZFP207-Halo fusion (primers are described in table S4). Zfp207-Halo plasmid was linearized, *in vitro* transcribed (MegaScript T7 Transcription kit, Thermo Fisher #AM1333) and *in vitro* translated (Wheat Germ *in vitro* translation kit, Promega #L4380). The *in vitro* translated Zfp207-Halo fusion product was incubated with total RNA from mouse ESCs. After this binding reaction, the RBP-RNA complex was purified and the bound RNA eluted for subsequent Illumina library preparation (NEXTFLEX small RNA library preparation kit v3, PerkinElmer #NOVA-5132-06) and paired-end (40×43 cycles) sequenced on an Illumina NextSeq 500. Sequencing reads were trimmed, adapters and ribosomal RNA reads removed and aligned against the reference genome GRCm38.p6 retrieved from GENCODE (https://www.gencodegenes.org/mouse/) using HISAT2 v2.1.0 with default parameters. Only uniquely mapping reads were further considered. PCR duplicate reads were removed. The libraries of the two replicates, HaloTag and input were read count normalized. Binding events (peaks) were called by using MACS2 v2.2.6 (https://github.com/macs3-project/MACS/wiki/Advanced:-Call-peaks-using-MACS2-subcommands). Genomic regions enriched after subtraction of Halo and Input signals are kept and further filtered at a FDR ≤ 0.01, a fold change (FC) enrichment FC≥3 (Halo) and FC≥2 (Input) and RPM≥2 in both replicates. Only consensus peaks were considered further. Genomic features (R Genomic Features package (v1.36.4) and R Genomic Ranges package (v1.36.1)) extracted and pathway enrichment was conducted using R ClusterProfiler (v3.12.0) to compute the significance of the enrichments and org.Mm.eg.db (v3.8.2) to retrieve Entrez IDs required by ClusterProfiler using the entire murine transcriptome as a background model. Each peak has a reported Binding Score computed as the base 2 logarithm of the product between the log^2^RPM and the mean enrichment over the two negative controls.

### Subcellular RNA fractionation

2 million cells were harvested and washed twice with PBS. After centrifugation for 10 minutes at 3,000 rpm, the pellets were resuspended in 2 packed volumes of cytoplasmic lysis buffer (10 mM Tris HCl pH 7.5, 0.15% NP-40, 150 mM NaCl supplemented with proteinase and RNase inhibitors) and mixed well by pipetting up and down 15 times. The mix was then incubated on ice for 10 minutes and 1.5 times volumes of chilled sucrose buffer (10 mM Tris HCl pH 7.5, 150 mM NaCl, 24% sucrose) were added to the lysate. The lysate was then centrifuged at 4 °C, 13,000 rpm for 3 minutes, and the supernatant was collected as cytoplasmic fraction. The nuclear pellet was resuspended in 2 packed volumes of cytoplasmic lysis buffer without NP-40 and mixed with 1.5 times volumes of chilled sucrose buffer followed by centrifugation at 13000 rpm 5 minutes at 4°C. The washed nuclear fraction was resuspended with chilled glycerol buffer (20 mM Tris HCL pH 7.9, 75 mM NaCl, 0.5 mM EDTA and 50% glycerol) followed by addition of equal volume of cold nuclei lysis buffer (10 mM HEPES pH 7.6, 1 mM DTT, 7.5 mM MgCl^2^, 0.2 mM EDTA, 0.3 M NaCl, 1 M Urea, 1% NP-40 supplemented with proteinase and RNase inhibitor). The mixture was vortexed twice for 5 seconds each and incubated on ice for 2 minutes followed by centrifugation at high-speed for 5 minutes. The supernatant was collected as a nuclear fraction. Total RNA from cytosolic and nuclear fraction were extracted using the RNeasy Mini Kit (Qiagen).

### RNA FISH

Cells were grown on microscope coverslips placed in tissue culture dishes. After washing twice with PBS, cells were fixed in 4% paraformaldehyde (Sigma-Aldrich) and permeabilized with 70% Ethanol for two hours at 4 °C. Following permeabilization, cells were washed twice with 25% of RNA wash buffer (25% formamide, 2X SSC) for 5 minutes at RT. Cells were stained with hybridization mix (10% dextran sulfate, 25% formamide, 2 x SSC, 1 mg/ml *E. coli* tRNA, 0.02% BSA, 10 mM vanadyl-ribonucleoside complex) and 2 µM 5’Cys3-oligo dT(50) probe for 24 hours at 30 °C. The following day, coverslips were washed with 25% of RNA wash buffer for 60 minutes at 30°C and incubated with 20 ng/ml DAPI (RNAse-free) in 25% RNA wash 30 minutes at 30°C. The coverslips were then rinsed with 2 x SSC buffer followed by RNA equilibration buffer (10 mM Tris HCl pH 7.5, 2x SSC and 0.4% glucose). Thereafter, coverslips were mounted using anti-bleach buffer (10 mM trolox, 37 ng/μl glucose oxidase, catalase in RNA equilibration buffer). Images were acquired using a Nikon microscope (ZEN lite software) and analyzed using ImageJ. For quantifying nuclear poly(A)^+^ RNA signals (N), images containing at least 30 nuclei were only considered. DAPI staining was masked and overlay on top of the 5’Cys3-oligo d(T) signal, after that intensity of pixel was measured using analyze particles. Then, the total FISH signal was measured as background fluorescence using a threshold setting. And the cytosolic signal was measured as subtracting the nuclear signal from the total signal. All the images were exposed to same magnitude of gain.

### Statistical analysis

Data are shown as mean ± SEM. GraphPad Prism version 8.0.0 for Windows was used to perform the statistical analysis (GraphPad Software, La Jolla California USA, www.graphpad.com). The significance was determined using student’s *t*-test, the ordinary one-way ANOVA and ratio paired *t*-test where indicated including correction for multiple comparison. Probability values of * p < 0.05, ** p < 0.001, *** p< 0.0001 were considered as statistically significant.

## General

We would like to thank Aguilo Lab members for useful discussion, the Biochemical Imaging Centre Umeå for assistance with the imaging processing, and David Muñoz Forcada for help with the figure’s presentation.

## Author contributions

F.A. conceived and designed the study. S.M., D.P.B. D.M.S and P.G. performed experiments, I.A. and C.K. performed RAP-seq analysis, A.C.R., D.Z. and D.F.L performed bioinformatics analysis. F.A. wrote the manuscript. All authors reviewed and edited the manuscript.

## Data and materials availability

All next generation sequencing data will can be publicly accessed in ArrayExpress webserver (E-MTAB-10108 and E-MTAB-10113).

