## Supplemental information for "ZFP207 *controls pluripotency by multiple post-transcriptional mechanisms*"

**This PDF file includes:**

Figs. S1 to S5

**Other Supplementary Materials for this manuscript include the following:**

Tables S1 to S4. See separate excel Files.

**Fig. S1.**

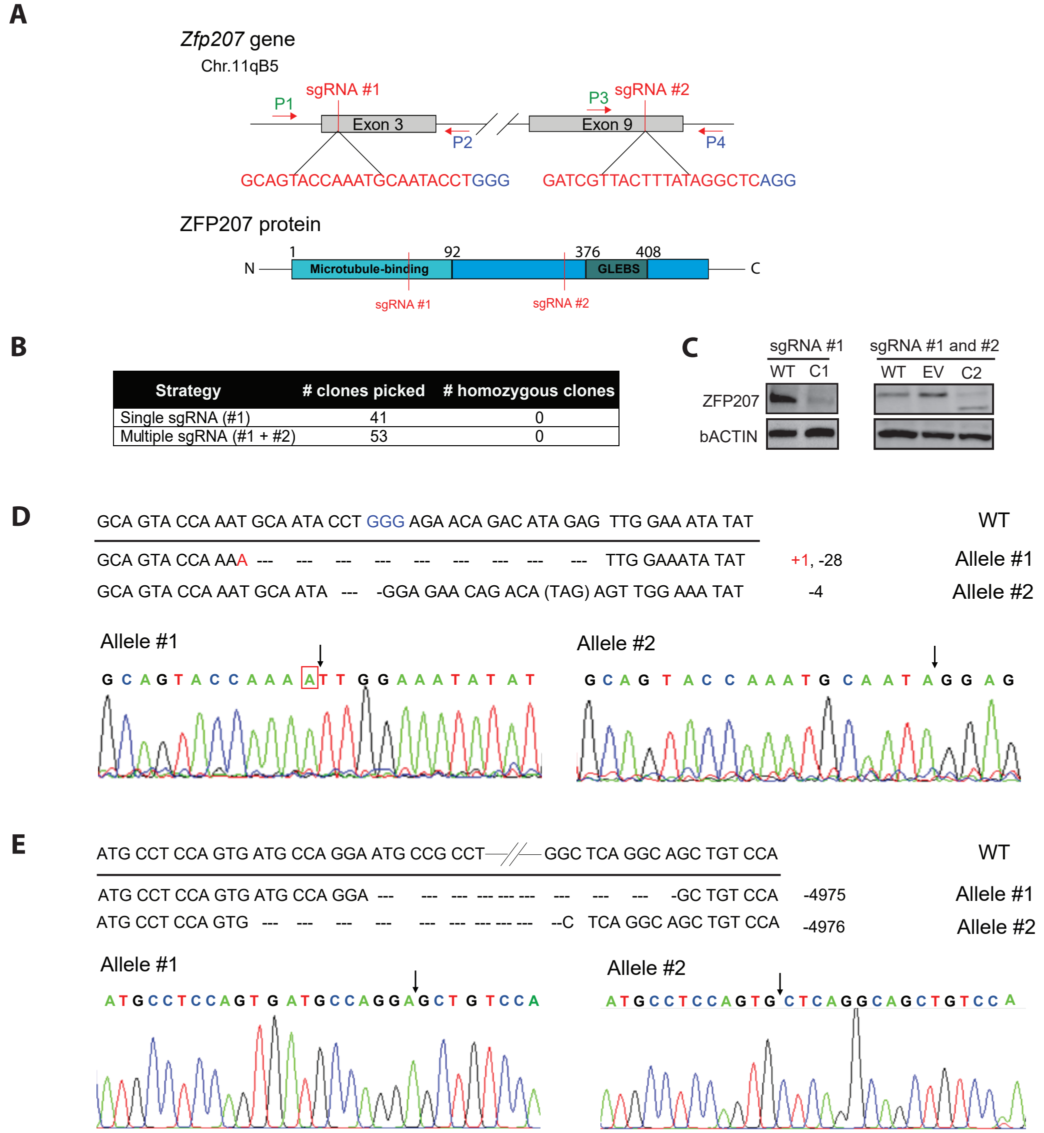


| Strategy | # clones picked | # homozygous clones |
|---|---|---|
| Single sgRNA (#1) | 41 | 0 |
| Multiple sgRNA (#1 + #2) | 53 | 0 |

**Figure S1. Generation of *Zfp207* KO mouse ESC lines, related to Figure 1.**

(**A**) Schematic diagram of sgRNA1 and sgRNA2 targeting the exons 3 and 9 of *Zfp207* gene, respectively (top) and corresponding sites of its encoded protein (bottom). The sgRNAs and PAM sequences are marked in red and blue, respectively. (**B**) Detailed information on the strategies used for KO generation, number of clones picked, and the number of homozygous clones obtained. (**C**) WB analysis of selected clones from two different strategies (sgRNA #1 and a combination of sgRNA #1 and #2) used to evaluate ZFP207 expression. β-ACTIN was used as a loading control. WT: Wild-type mouse ESC; C1: clone 1; EV: empty vector (px459); and C2: clone 2. (**D**, **E**) Sanger sequencing analysis (upper panel) and chromatogram (lower panel) of selected clones C1 and C2. Insertion is indicated in red while deletions are represented in dashes and in black arrows in the chromatogram.

**Fig. S2.**

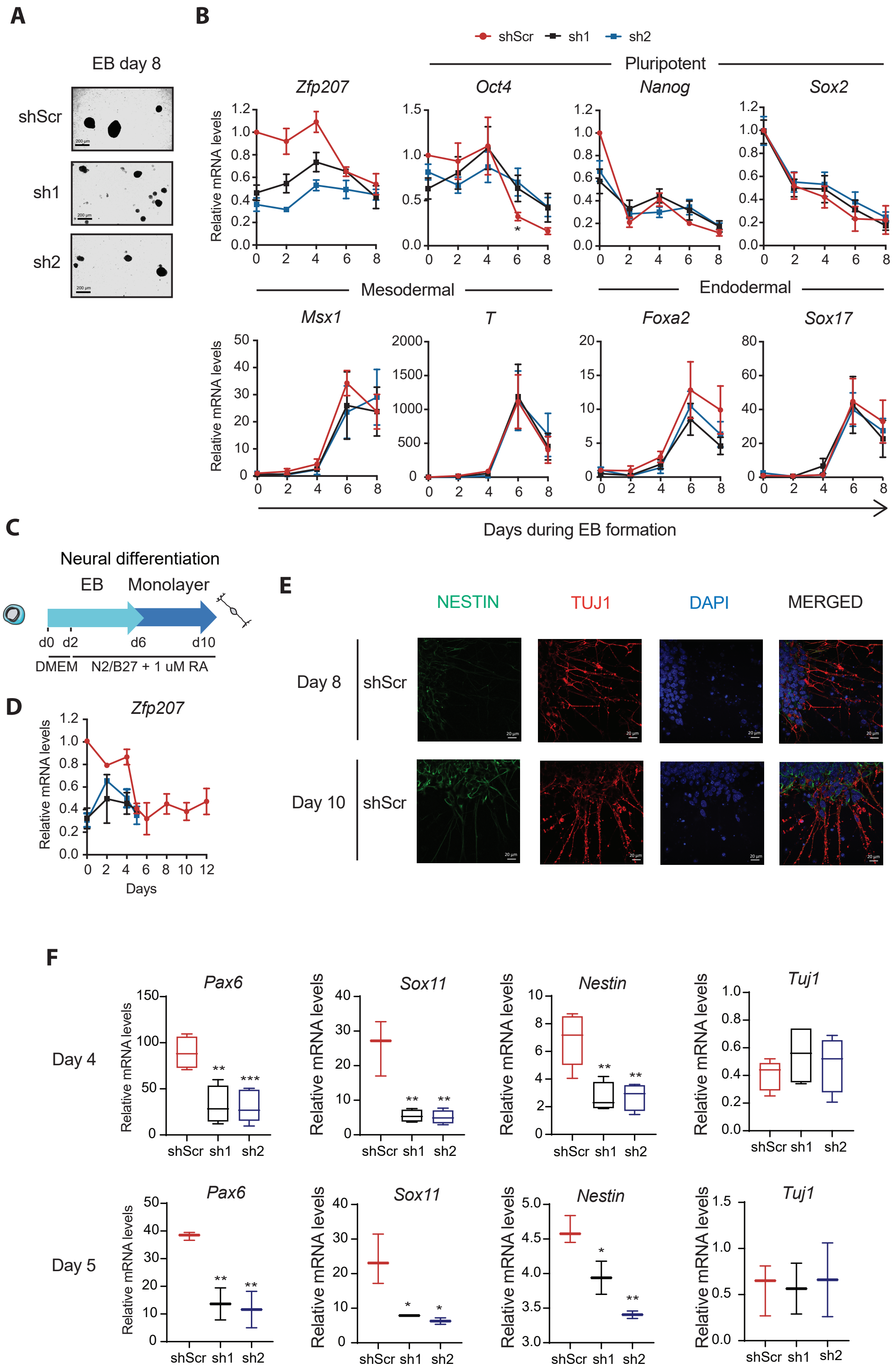


*Figure legend in the next page.*

**Figure S2. Ablation of *Zfp207* disrupts differentiation, related to Figure 2.**

(**A**) Representative bright-field images (4x magnification) of embryoid bodies (EB) generated from control (shScr), and *Zfp207*-depleted ESCs (sh1 and sh2) at day 8 of differentiation. Scale bars, 200 μM. (**B**) Gene expression analysis of *Zfp207,* pluripotency markers (*Oct4, Nanog* and *Sox2*), mesoderm-associated genes (*Msx1* and *T*) and endoderm markers (*FoxA2* and *Sox17*) in shScr, sh1 and sh2 ESCs along the course of EB-mediated differentiation. mRNA levels are normalized to *β-Actin* and represented as mean ± SEM relative to the expression of shScr at day 0; n =3, * $p < 0.05$. A ratio paired Student's *t*-test is used for statistical analysis. (**C**) Schematic depiction for the neuroectodermal-directed differentiation. DMEM/F12 supplemented with N2B27and retinoic acid (RA) was added after two days of EB culture. (**D**) RT-qPCR analysis of *Zfp207* in shScr, sh1 and sh2 along the time-course of neuroectodermal differentiation. Data points for sh1 and sh2 were profiled up to day 5 because the cells died. *Zfp207* is normalized to *β-Actin* and represented as mean ± SEM relative to the expression of shScr at day 0; n = 3. (**E**) Immunostaining of NESTIN (green) and TUJ1 (red) of neural progenitors generated from shScr on day 8 and 10 of the neuroectodermal differentiation. Nuclei were counterstained with DAPI. Scale bar, 20 μM. (**F**) RT-qPCR of the neural-associated markers (*Pax6, Sox11, Nestin, and Tuj1*) at the indicated time points of neural directed differentiation in shScr and *Zfp207*-depleted ESCs (sh1 and sh2). mRNA levels are normalized to *β-Actin* and represented as mean ± SEM relative to the expression of shScr at day 0; n =3, * $p < 0.05$, ** $p < 0.001$, *** $p < 0.0001$. An unpaired Student's *t*-test is used for statistical analysis.

**Fig. S3.**

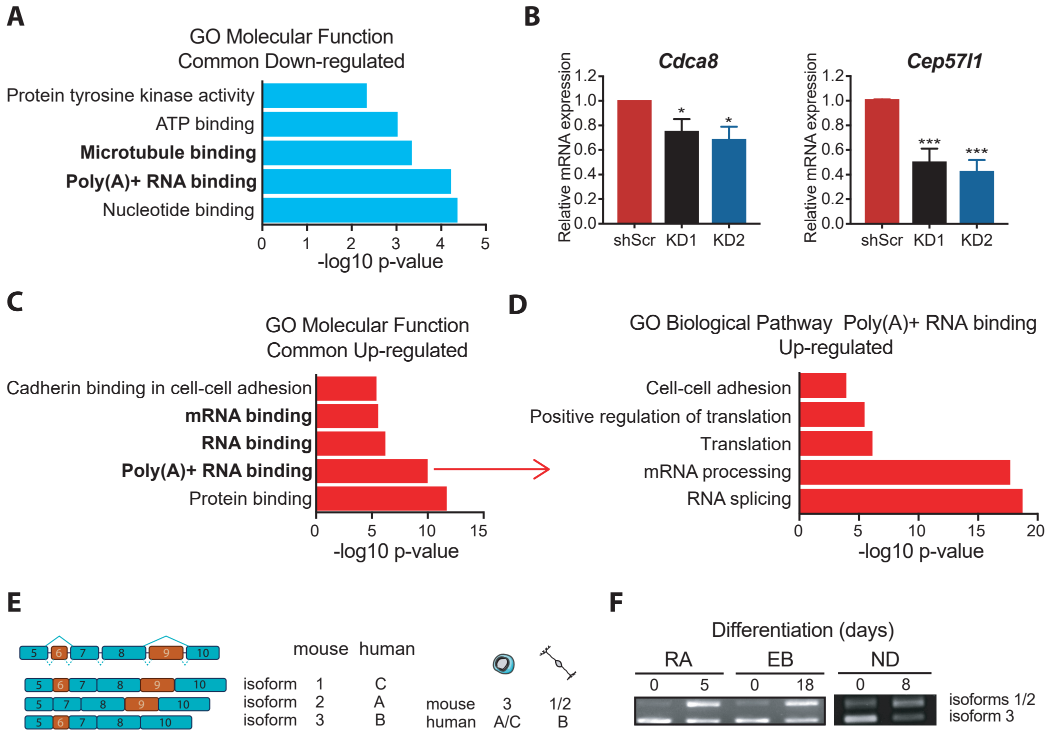


**Figure S3. Identification of ZFP207 targets, related to Figure 4.**

(**A**) Gene ontology (GO) analysis of molecular functions associated with common down-regulated genes in *Zfp207*-depleted ESCs (sh1 and sh2) compared to shScr. (**B**) RT-qPCR analysis of mitotic sister chromatid segregation related genes (*Cdca8* and *Cep57l1*) in shScr, sh1 and sh2 ESCs. mRNA levels are normalized to *β-Actin* and represented as mean ± SEM relative to the expression of shScr; n =3, * $p < 0.05$, *** $p < 0.0001$. An unpaired Student's *t*-test is used for statistical analysis. (**C**) GO analysis of molecular functions associated with common up-regulated genes and (**D**) of biological processes associated with poly(A)+ RNA binding up-regulated genes in *Zfp207*-depleted ESCs (sh1 and sh2) compared to shScr. **(E)** Schematic representation of three different AS forms of *Zfp207* in mouse and human, respectively (left panel). And the differential switch of the splice forms between ESC state and neural-directed differentiation in mouse and human (right panel). (**F**) Representative RT-PCR analysis of the AS forms of *Zfp207* at the indicated time points of retinoic acid (RA) -induced differentiation, embryoid body (EB) generation and neuroectodermal (NE) differentiation of mouse ESCs.

**Fig. S4.**

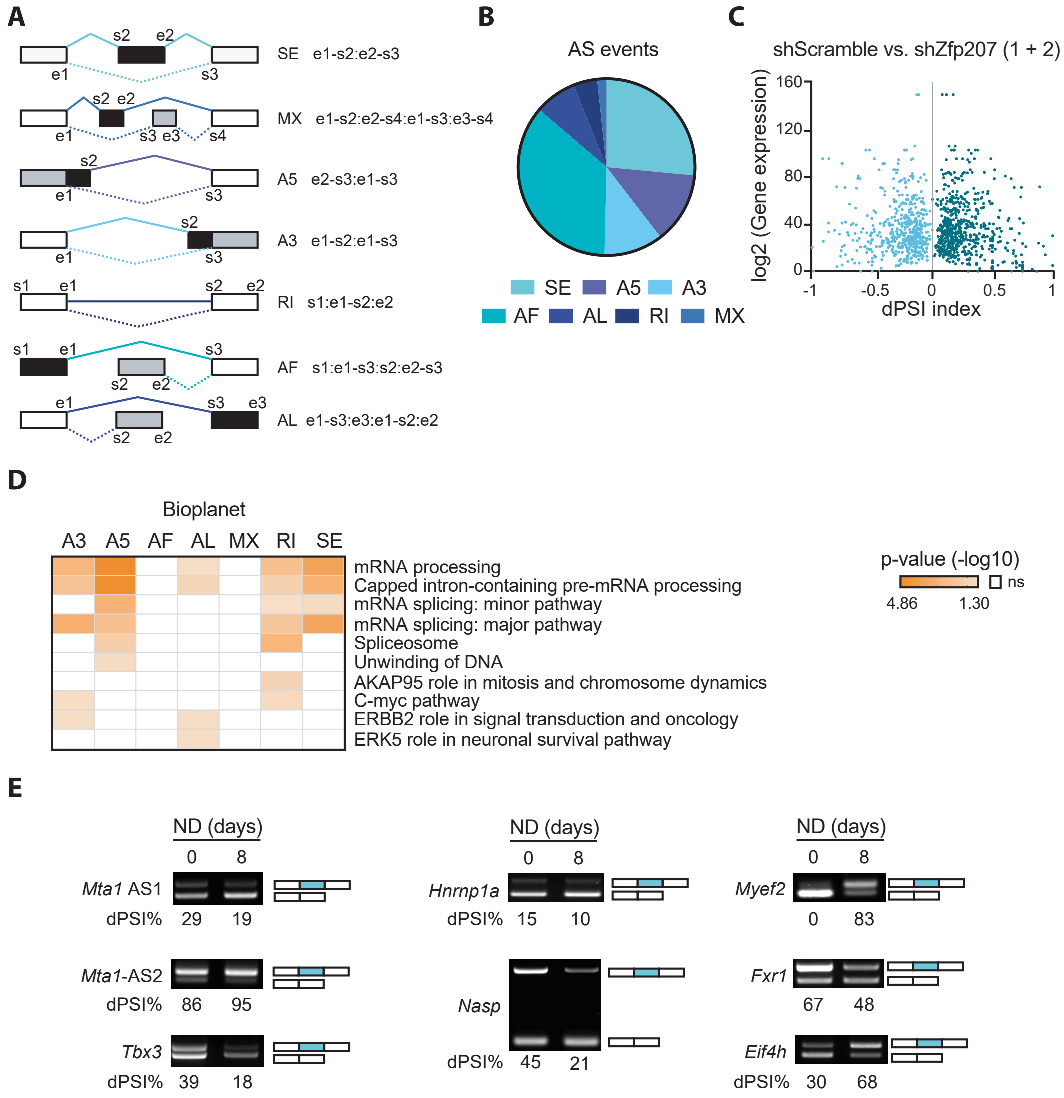


**Figure S4. Alternative splicing switches in *Zfp207* knockdown ESCs are prevalent in differentiated cells, related to Figure 5.**

(**A**) Schema depicting the events generated by the software SUPPA2 with the specific coordinates, including start (s) and end (e). The form of the AS event is depicted in black. (**B**) Pie chart showing the AS events types distribution. (**C**) Volcano plot depicting the correlation between gene expression levels and dPSI index resulted from RNA-seq data analysis in control (scShr) and *Zfp207* depleted mouse ESCs (sh1 + sh2). Gene expression levels are presented as log2 transformed values. dPSI index range is between -1 and +1. (**D**) Gene ontology enrichment analysis with Bioplanet software for genes undergoing different splicing events upon *Zfp207* depletion (sh1 and sh2). (**E**) Representative RT-PCR analysis of AS events upon neural differentiation in mouse ESCs for *Mta1* (AS1 and AS2), *Tbx3*, *Hnrnp1a, Nasp*, *Myef2*, *Fxr1,* and *Eif4H*. The structure of each isoform is indicated (not to scale). Alternative exons are blue. The percent spliced in (PSI) was quantified for each condition.

**Fig. S5.**

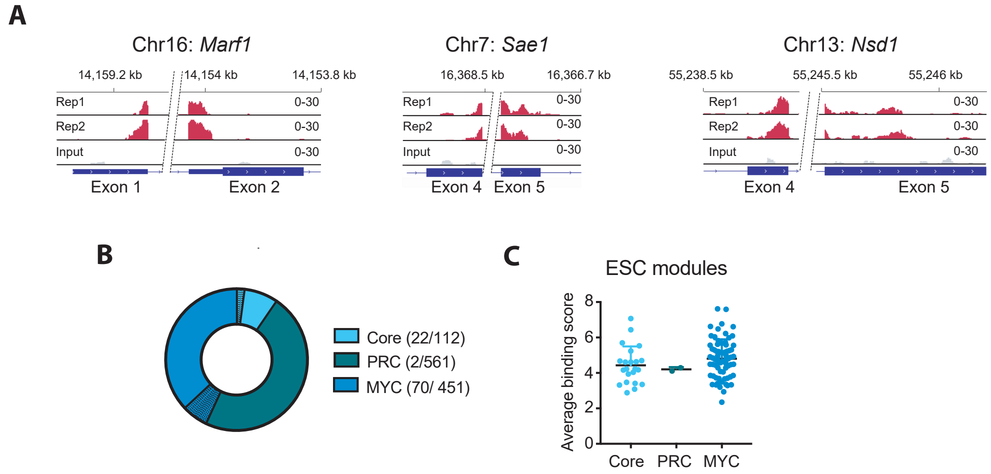


**Figure S5. ZFP207 binding profile to mRNAs, related to Figure 6.**

(**A**) RAP-seq binding profile for ZFP207 at *Marf1*, *Sae1*, and *Nsd1* genes. Values on the y axis represent a sequencing depth normalized and HaloTag subtracted read-count. Both replicates (Rep1 and Rep2; red) and the input (grey) are shown. (**B**) Pie chart depicting ZFP207 binding sites (black lines) at the Core, PRC and Myc modules and (**C**) its average binding score.

Table S1: RNA-Seq. See separate excel File, Table S1.xlsx

Table S2: AS genes. See separate excel File, Table S2.xlsx

Table S3, RAP-seq for ZFP207 immunoprecipitated from mouse ESCs. See separate excel File, Table S3.xlsx

Table S4, related to experimental procedures: primers used for gene expression analysis, AS validation, CRISPR-Cas9, and cloning. See separate excel File, Table S4.xlsx
